# Sclerotome is compartmentalized by parallel Shh and Bmp signaling downstream of CaMKII

**DOI:** 10.1101/2023.07.21.550086

**Authors:** Sarah C. Rothschild, Richard H. Row, Benjamin L. Martin, Wilson K. Clements

**Affiliations:** Life Sciences, Virginia Commonwealth University, Richmond, VA 23284, USA; Department of Biochemistry and Cell Biology, Stony Brook University, Stony Brook, NY, 11794-5215, USA; Department of Hematology, St. Jude Children’s Research Hospital, Memphis, TN, 38105, USA

**Keywords:** CaMKII, Shh, Bmp, sclerotome, zebrafish, somite

## Abstract

The sclerotome in vertebrates comprises an embryonic population of cellular progenitors that give rise to diverse adult tissues including the axial skeleton, ribs, intervertebral discs, connective tissue, and vascular smooth muscle. In the thorax, this cell population arises in the ventromedial region of each of the segmented tissue blocks known as somites. How and when sclerotome adult tissue fates are specified and how the gene signatures that predate those fates are regulated has not been well studied. We have identified a previously unknown role for Ca^2+^/calmodulin-dependent protein kinase II (CaMKII) in regulating sclerotome patterning in zebrafish. Mechanistically, CaMKII regulates the activity of parallel signaling inputs that pattern sclerotome gene expression. In one downstream arm, CaMKII regulates distribution of the established sclerotome-inductive morphogen sonic hedgehog (Shh), and thus Shh-dependent sclerotome genes. In the second downstream arm, we show a previously unappreciated inductive requirement for Bmp signaling, where CaMKII activates expression of *bmp4* and consequently Bmp activity. Bmp activates expression of a second subset of stereotypical sclerotome genes, while simultaneously repressing Shh-dependent markers. Our work demonstrates that CaMKII promotes parallel Bmp and Shh signaling as a mechanism to first promote global sclerotome specification, and that these pathways subsequently regionally activate and refine discrete compartmental genetic programs. Our work establishes how the earliest unique gene signatures that likely drive distinct cell behaviors and adult fates arise within the sclerotome.

## Introduction

Somites are epithelialized metameric structures in vertebrates derived from mesoderm found adjacent to the neural tube and notochord (Christ et al., 2004; Monsoro-Burq, 2005; Stickney et al., 2000). During somite maturation, the somite is compartmentalized into regions containing precursors with distinct adult tissue fates. The ventromedial compartment of the somite—known as the sclerotome—undergoes an epithelial to mesenchymal transition and migrates to sites of adult differentiation as skeletal and connective tissue components of the vertebral column, ribs, sternum, vascular smooth muscle cells, and other tissues (Christ et al., 2004; Kato and Aoyama, 1998; Kozhemyakina et al., 2015; Monsoro-Burq, 2005; Stickney et al., 2000; Stratman et al., 2017).

Although sclerotome-derived cells contribute to diverse adult tissues, the time at which these fate potentials are originally specified and the distinct gene signatures that drive unique migration, maturation, and differentiation programs are poorly understood. Examination of gene signatures within the early sclerotome reveals non-uniform gene expression, with the most obvious compartmentalization falling between anterior and posterior regions (Christ et al., 2004; Hughes et al., 2009; Monsoro-Burq, 2005). These unique gene signatures reveal the earliest programming differences within the sclerotome compartment, but little is known about the mechanisms that establish them.

The most well characterized input required for sclerotome specification is sonic hedgehog (SHH) signaling from the notochord and floor plate (Chiang et al., 1996; Fan and Tessier-Lavigne, 1994; Johnson et al., 1994; Teillet et al., 1998). Inhibition of SHH via loss of the signal-activating receptor, smoothened (Smo), or loss of the glioma associated oncogene (Gli) family SHH-dependent transcription factors, Gli1, Gli2, Gli3, causes a loss of key sclerotome markers and consequent defects in axial skeleton and other tissues (Buttitta et al., 2003; Cairns et al., 2008; Chiang et al., 1996; Zhang et al., 2001). SHH secretion and signaling directs specification of the sclerotome through activation of paired-box containing transcription factors PAX1 and PAX9 (Peters et al., 1999). *Pax1* mouse mutants develop malformed vertebral column, sternum, and scapula, (Wilm et al., 1998). *Pax1;Pax9* double mutants fail to form sclerotome and do not generate the axial skeleton, thus demonstrating even more severe defects in sclerotome-derived adult tissues than *Pax1* mutants alone (Peters et al., 1999). Thus, SHH signaling and downstream transcriptional programs are a key component of sclerotome maturation and tissue differentiation.

Although sclerotome formation has been shown to be induced and patterned by nearby tissues including the notochord, floorplate, and ectoderm (Monsoro-Burq, 2005), few responsible signaling pathways outside of SHH have been identified or characterized. We previously showed that β-catenin-independent (non-canonical) Wnt signaling downstream of Wnt16 was required for optimal sclerotome gene expression in zebrafish (Clements et al., 2011). In the absence of Wnt16 signaling, there are significant downregulations in expression of diverse sclerotome markers and perturbations in regional expression and maturation.

A negative regulatory role for BMP signaling in sclerotome formation has been inferred from the finding that secreted BMP antagonists, notably noggin and follistatin, are required for normal early expression of the sclerotome markers *Pax1* and *Pax9* in mouse (McMahon et al., 1998; Stafford et al., 2011; Stafford et al., 2014). These findings point to the idea that BMP signaling represses sclerotome and that relief from BMP signaling by expression of secreted BMP inhibitors is critical to allow sclerotome induction. Most current models of sclerotome formation thus posit BMP signaling as antagonistic to early sclerotome development.

Although Ca^2+^ signaling has not yet been implicated in sclerotome specification, it is involved in overall somite formation and maturation. During the segmentation period, calcium transients are observed within the somites of diverse vertebrates (Webb and Miller, 2006). Inhibition of Ca^2+^ signaling, using chelators or antagonists inhibits somite segmentation in chick and zebrafish embryos (Chernoff and Hilfer, 1982; Webb and Miller, 2006).

A key molecule important in transducing Ca^2+^-dependent cellular responses is Ca^2+^/calmodulin-dependent protein kinase (CaMKII; Tombes et al., 2003). CaMKII is a serine/threonine protein kinase that is encoded by four genes (α, β, γ, δ) in humans (Tombes et al., 2003). There are seven zebrafish CaMKII orthologs (α1, β1, β2, γ1, γ2, δ1, δ2) that are 90-96% identical to their human counterparts (Rothschild et al., 2009; Rothschild et al., 2007). Inhibition of CaMKII results in defects in somite-derived adult tissues, for example in muscle cells it impairs myosin thick-filament assembly and alters the calcium-induced calcium release necessary for skeletal muscle contraction (Anderson et al., 1998; Ferrari and Spitzer, 1999; Wehrens, 2011). CaMKII is also necessary for chondrogenesis and has been linked to osteoarthritis development in adults (Shimazaki et al., 2006). Thus Ca^2+^ and its downstream effector CaMKII have been implicated in global somite patterning but a role in sclerotome development has not yet been investigated.

Here we identify a previously unknown early requirement for γ1 CaMKII in sclerotome specification in zebrafish embryos. Mechanistically, γ1 CaMKII regulates distribution of Shh and regulates its later transcriptional expression. In parallel, it is required for expression of *bmp4*, with disruption leading to loss of downstream Bmp signaling. Unexpectedly, we show a role for Bmp signaling in positively regulating a specific subset of sclerotome genes distinct from the established sclerotome markers it antagonizes, *pax1a* and *pax9*. Our results demonstrate that discrete subsets of sclerotome genes are differentially regulated by these parallel signaling pathways, and that permissive and antagonistic interactions between the pathways provide a mechanism for initiation and refinement of early sclerotome subcompartments. Our study thus identifies γ1 CaMKII as a pivotal signaling node upstream of Bmp and Shh signaling, which in turn subsequently drive establishment of anterior and posterior gene signatures within the sclerotome.

## Results

### *Camk2g1* is required for sclerotome specification

In zebrafish, genetic markers of sclerotome development, including *pax1a*, *pax9*, *foxc1a*, *twist1b*, *twist2*, and *snai2*, initiate and are expressed in ventromedial cells of the somite from around 16-18 hours post fertilization (hpf). Consistent with a potential role in sclerotome development, *camk2g1* mRNA is expressed in ventral cells of the somite at 12-26 hpf (Fig. S1 A-C, arrows) as well as the notochord during early somite stages. We verified CaMKII protein expression in this region using an antibody that recognizes both zebrafish γ1 and δ2 CaMKII polypeptides, revealing protein in the notochord, spinal cord, and spinal cord cell bodies at 24 hpf (Fig. S1 E). Of the two polypeptides recognized by the antibody, only *camk2g1* message is expressed in the notochord and neural tube at this time point, so protein expression is most likely specific to γ1 CaMKII (Rothschild et al., 2007). We conclude that *camk2g1* is expressed in tissues that contribute to and regulate sclerotome specification, including the ventral somite mesoderm and notochord.

To evaluate the role of *camk2g1* in sclerotome specification in zebrafish embryos we employed parallel antisense and genetic loss of function approaches. Embryos were injected with a previously validated translation-blocking antisense morpholino (MO) (Francescatto et al., 2010; Rothschild et al., 2011; Rothschild et al., 2013) or Cas9 protein and four gRNAs targeting *camk2g1* (Rothschild et al., 2020). Injection of morpholino caused a dramatic loss of CaMKII protein in both the spinal cord cell bodies and notochord (Fig. S1 F), confirming its efficacy. In parallel, we employed CRISPR-targeted genetic deletion of *camk2g1*, which we previously confirmed and which overcomes genetic compensation by *camk2g2* observed in TALEN-directed stable mutants (Rothschild et al., 2020). Knockdown of γ1 CaMKII by both means reduced expression of a panel of known sclerotome markers including *foxc1a*, *twist1b*, *twist2*, *snai2* and *pax9*, while *pax1a* expression was diminished and appeared disorganized at 24 hpf (Fig. 1, Table S1). *Foxc1a*, *twist1b*, and *snai2* are preferentially located in anterior sclerotome, while *pax1a*, *pax9*, and *twist2* are found in posterior sclerotome. Our results indicate that *camk2g1* is expressed in sclerotome and nearby inductive tissues, and that it is required for sclerotome development.

**Fig. 1.**
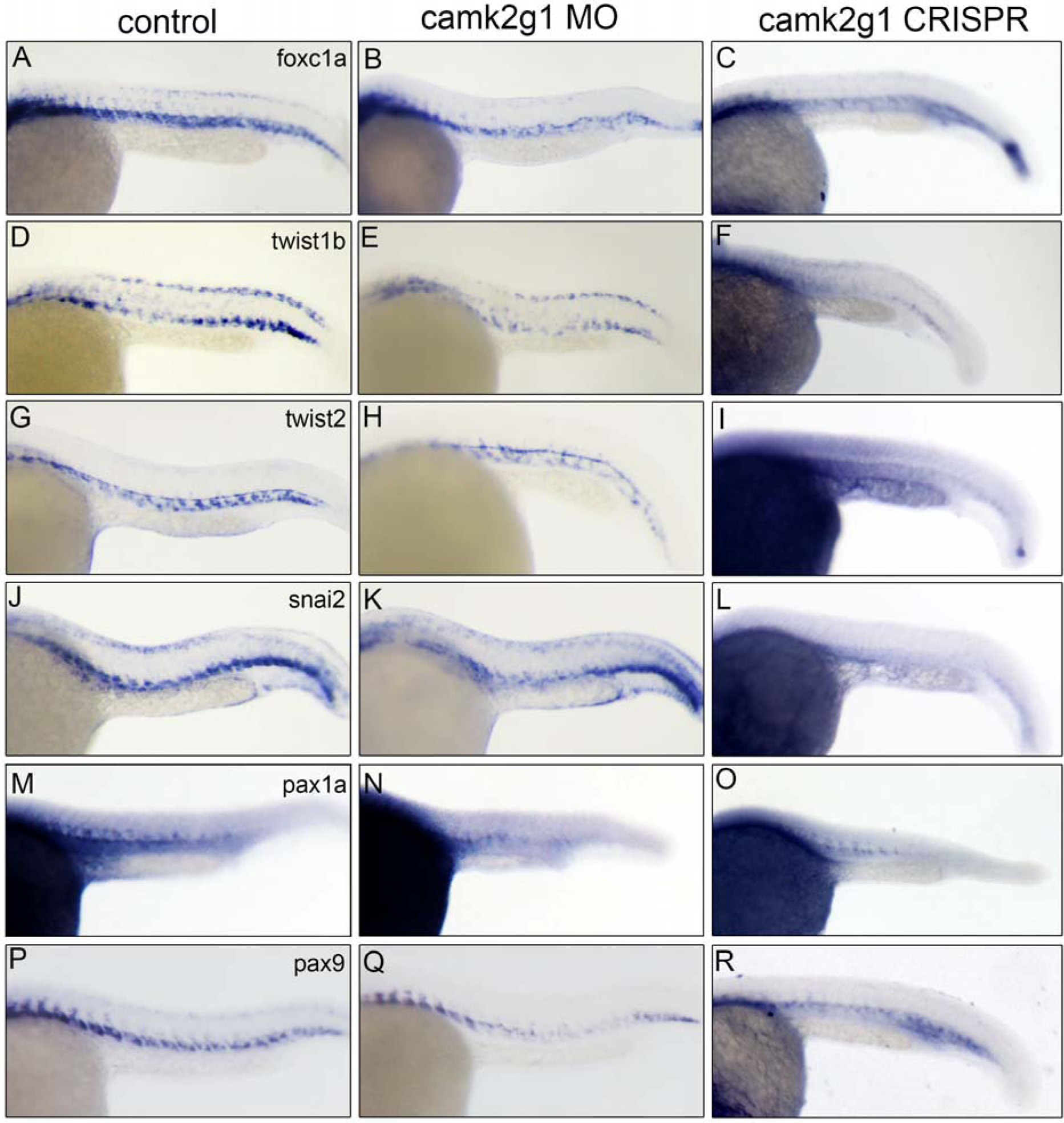
γ1 CaMKII loss of function leads to decreased sclerotome gene expression. Sclerotome development was analyzed by marker expression in control (left), *camk2g1* MO-injected embryos (middle), and *camk2g1* CRISPR/Cas9-injected embryos (right) at 24 hpf. *Foxc1a* (A, B), *twist1b* (C, D), *twist2* (E, F), *snai2* (G, H), *pax1a* (I, J) and *pax9* (K, L) are reduced in *camk2g1* morphants compared to controls. Lateral views, anterior left, dorsal up.

### Inhibition of Shh and Bmp signaling inhibits sclerotome development

Sclerotome development is regulated by signals from nearby tissues (Stickney et al., 2000). Shh secretion from the notochord initiates expression of the sclerotome markers *pax1a* and *pax9* (Abbott and Ducibella, 2001; Buttitta et al., 2003; Peters et al., 1999). To better understand genes regulated by Shh, we treated embryos with a pharmacological inhibitor of Shh signaling, the smoothened antagonist cyclopamine. Expression of the anterior sclerotome markers *pax1a* and *pax9* was reduced, as has been observed previously (Fig. 2 A-D, Table S1; Arnold et al., 2015; Ma et al., 2018), and we additionally noted significant loss of *twist2* (Fig. 2 E-F, Table S1). In contrast, expression of *foxc1a, twist1b*, and *snai2* remained unchanged in cyclopamine-treated embryos (Fig. 2 G-L, Table S1), indicating that their expression is independent of Shh at this time point.

**Fig. 2.**
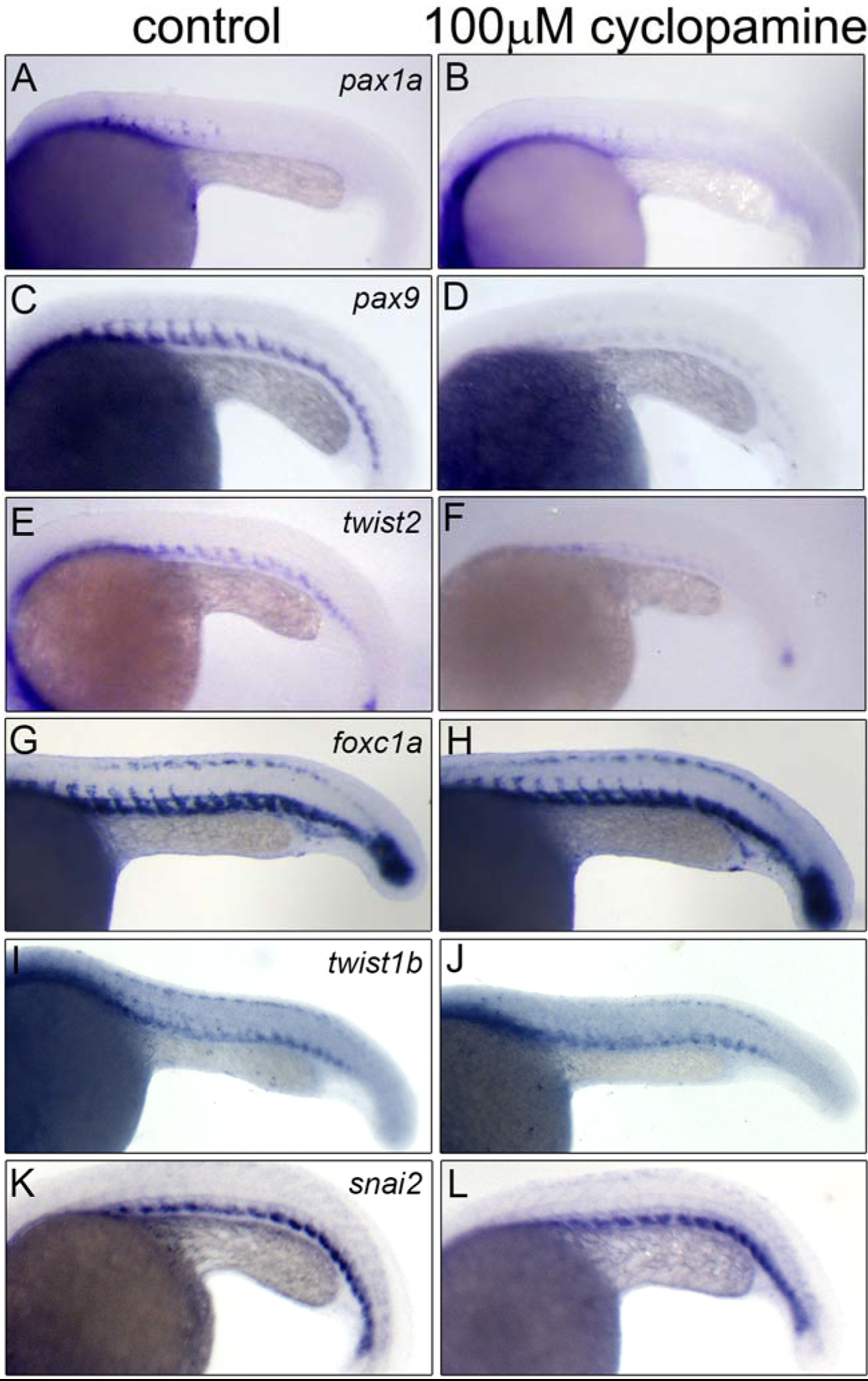
Inhibition of Shh signaling reduces expression of specific sclerotome genes. Embryos were treated with vehicle (left) or 100 μM cyclopamine (right) for 3 hrs, fixed, and analyzed for sclerotome marker expression at 22 hpf. *Pax9* (A, B), *pax1a* (C, D), and *twist2* (E, F) are diminished in cyclopamine-treated embryos. *Foxc1a* (G, H), *twist1b* (I, J), and *snai2* (K, L) are unaffected after cyclopamine treatment. Lateral views, anterior left, dorsal up.

The fact that not all sclerotome gene expression was altered by Shh inhibition indicated additional inductive regulatory inputs. Bmp pathway factors, including Bmp ligands and phospho-Smads-1,5,8/9 (pSmad-1,5,8/9), which transduce Bmp signaling, are found in the ventral mesoderm during the time window when sclerotome develops (Collery and Link, 2011; Dick et al., 1999; Laux et al., 2011). To determine if Bmp signaling might influence early sclerotome specification, we inhibited endogenous Bmp signaling using two separate parallel approaches: pharmacological inhibition of Bmp signaling and expression of a dominant negative Bmp receptor by heat shock using *hs:dnBmpr-GFP* transgenic fish (Hao et al., 2010; Pyati et al., 2006; Pyati et al., 2005). Embryos incubated with 10 μM DMH1, a Bmp antagonist, displayed greatly diminished expression of *foxc1a*, *twist1b*, and *snai2* (Fig. 3 A-F1, Table S1), but little or no alteration in *pax1a*, *pax9*, or *twist2* (Fig. 3 G-L, Table S1), genes that we previously observed to require Shh signaling. Similarly, ubiquitous expression of dnBmpr decreased *foxc1a* and *twist1b* expression in the sclerotome (Fig. S2). Because *Bmp4* is a known transcriptional target of Shh during stem cell regeneration (Zheng et al., 2013), we tested *bmp4* expression and Bmp signaling activity in a *BRE:EGFP* Bmp signaling reporter transgenic (Collery and Link, 2011) with or without cyclopamine. Expression of the *EGFP* message (a more sensitive and immediate readout of active Bmp signaling than fluorescence) in *BRE:EGFP* reporters was not altered in cyclopamine-treated embryos (Fig. S3 A, B). Interestingly, *bmp4* expression slightly increased after cyclopamine treatment (Fig. S3 C, D). Our results indicate that ventral somite Bmp expression and signaling are independent of Shh and must therefore be regulated by other factors. Thus, a subset of genes (*foxc1a*, *twist1b*, *snai2*) are regulated by Bmp and a second subset of genes (*pax1a*, *pax9*, *twist2*) are regulated by Shh. Our results demonstrate that Bmp and Shh signaling are independently and differentially required for expression of distinct sclerotome genes within the broader sclerotome in zebrafish embryos and lead to the activation of specific sclerotome gene signatures.

**Fig. 3.**
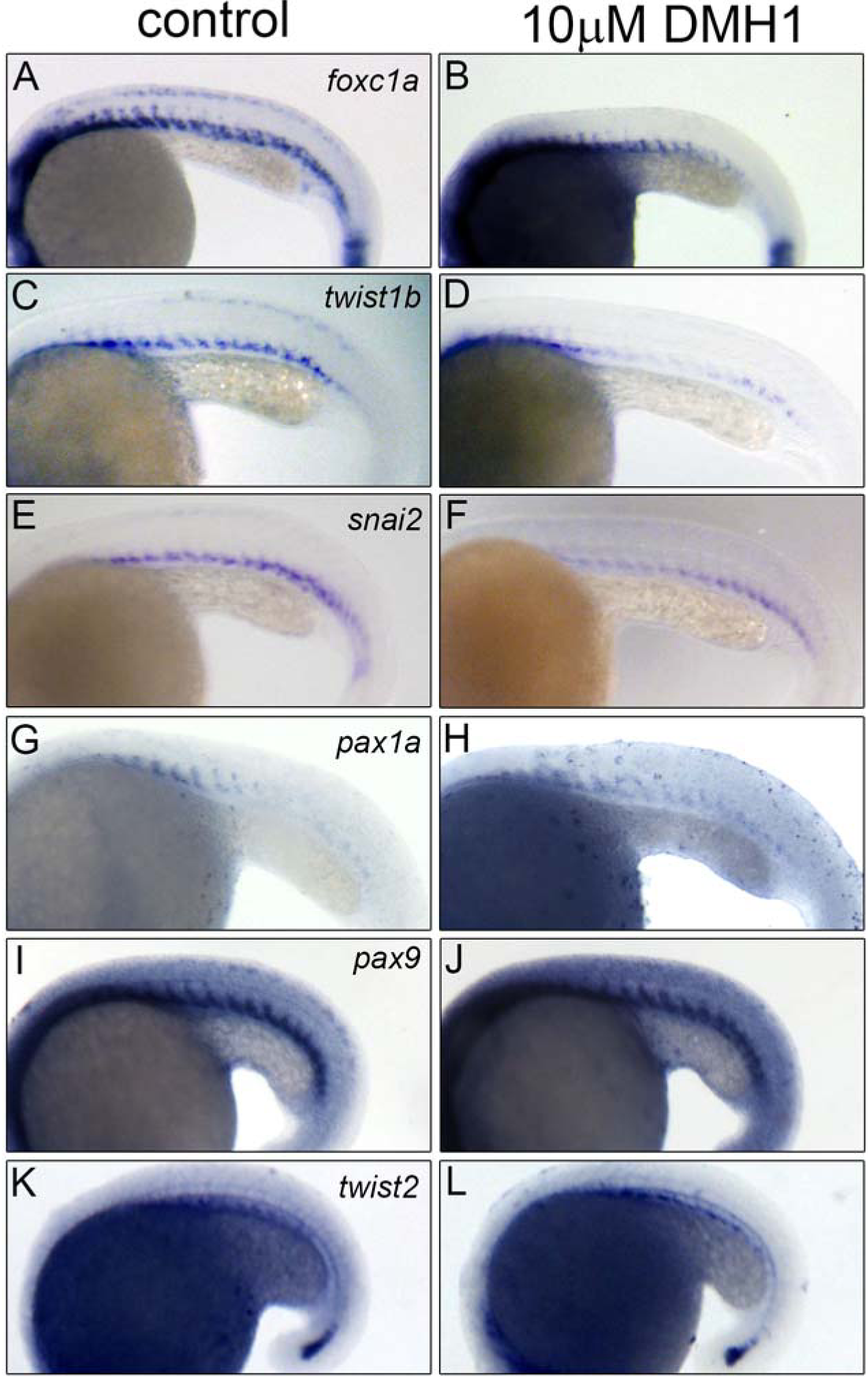
Inhibition of BMP signaling decreases expression of specific sclerotome genes. Embryos were treated with vehicle (left) or 10 μM DMH1 (right) for 2 hrs, fixed, and analyzed for sclerotome gene expression at 20 hpf (A-H) or 18 hpf (I-L). *Foxc1a* (A, B), *twist1b* (C, D), and *snai2* (E, F) are reduced in DMH1 treated embryos, while *pax1a* (G, H), *pax9* (I, J), and *twist2* (K, L) are expressed normally compared to controls. Lateral views, anterior left, dorsal up.

### Shh secretion and gradient formation is altered in *camk2g1* morphants

*Camk2g1* morphants have reduced expression of the Shh targets *pax9*, *pax1a*, and *twist2* in the sclerotome suggesting that Shh signaling might somehow be impaired. To better understand the basis of altered sclerotome gene expression in tissues neighboring the notochord and floorplate, we compared Shh secretion and gradient formation by immunofluorescence in untreated embryos or *camk2g1* morphants. Transverse sections of uninjected control embryos revealed robust Shh puncta throughout the somite, notably in the sclerotome compartment at 18 hpf (Fig. 4 A, B, B’). In contrast, *camk2g1* morphant embryos had reduced Shh puncta—especially in the ventral somite—compared to control embryos (Fig. 4 A, C, C’), indicating defects in Shh secretion and gradient formation. Our results reveal that γ1 CaMKII diminishes packaging and distribution of Shh at time points relevant to sclerotome specification.

**Fig. 4.**
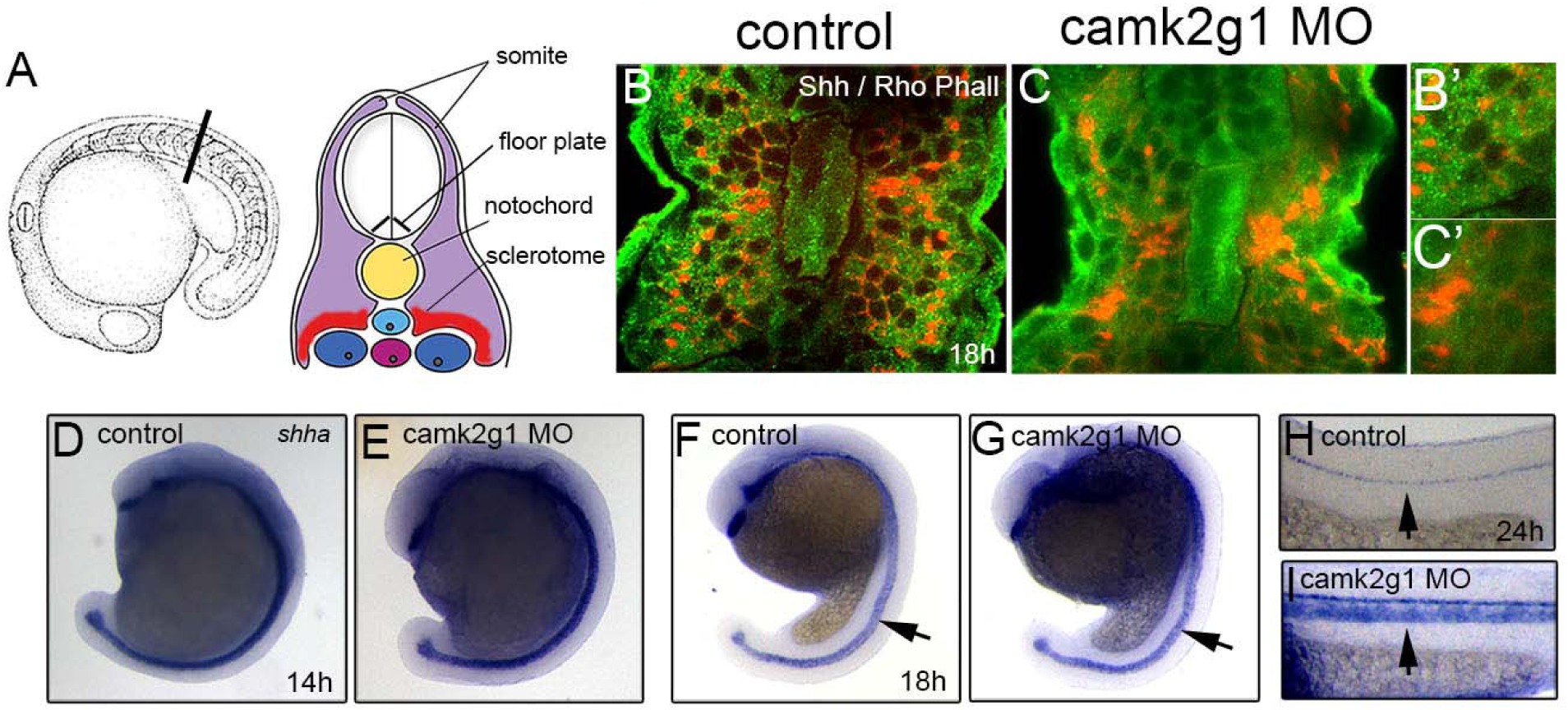
γ1 CaMKII knockdown inhibits Shh secretion and distribution, and alters *shha* message levels. (A) Schematic of embryo and regions presented in B, C. Transverse sections (dorsal up) showing the distribution of Shh protein detected by immunohistochemistry (green) in control (B) or *camk2g1* morphants (C) at 18 hpf reveal distinct Shh puncta in the ventral region of a representative uninjected embryo (B, magnified image B’). *Camk2g1* morphants lack ventral Shh puncta but have increased Shh in the notochord (C, magnified image C’). *Shha* mRNA expression in control (D, F, H) and *camk2g1* morphants (E, G, I) is normal at 14 hpf, but increased slightly at 18 hpf (F vs. G, arrows) and strongly 24 hpf in *camk2g1* morphants (H vs. I, arrows). D-G, lateral views, anterior up, dorsal right. H, I lateral trunk views, anterior left, dorsal up.

Shh signaling is subject to robust feedback mechanisms (Briscoe and Therond, 2013; Jeong and Epstein, 2003; Koudijs et al., 2008; Laufer et al., 1994; Lee et al., 2016; Niswander et al., 1994; Varjosalo and Taipale, 2008), so we wondered if incorrect secretion of Shh in *camk2g1* morphants might affect *shh* mRNA expression. Control and *camk2g1* morphant embryos had equivalent levels of *shha* transcript in the notochord and floor plate at 14 hpf; however, at 18 hpf *camk2g1* morphants had elevated *shha* expression in the notochord (Fig. 4 D-G), and at 24 hpf, when uninjected controls had mostly terminated *shha* expression in the notochord, *camk2g1* morphants still had robust expression of *shha* throughout (Fig. 4 H, I). Our data are consistent with a model where under normal circumstances, Shh secreted from the notochord upregulates a negative feedback mechanism that causes downregulation of *shha* in the notochord. When Shh secretion is impaired by loss of γ1 CaMKII, the negative feedback loop is interrupted and notochord downregulation of *shha* does not occur, leading to proglonged expression. Together, our results identify γ1 CaMKII as an important regulator of Shh secretion, gradient formation, and expression.

### Knockdown of γ1 CaMKII inhibits Bmp signaling

Bmp-inhibited animals, like *camk2g1* morphants, exhibit reduced *foxc1a*, *twist1b*, and *snai2* expression (Figs. 1, 3), suggesting that Bmp might mediate γ1 CaMKII effects. To explicitly test this possibility, we examined whether Bmp signaling activity was reduced during sclerotome maturation in Bmp reporter embryos (*BRE:EGFP*) that were either uninjected or injected with *camk2g1* morpholino. At 16 and 22 hpf, EGFP fluorescence was massively reduced in the tails of *camk2g1* morphants compared to controls (Fig. 5 A-F). Analysis of *EGFP* transcript levels by *in situ* in *camk2g1* morphant embryos at 18 and 22 hpf confirmed the decrease (Fig. 5 G-J). Like EGFP fluorescence, levels of the active form of Bmp signal transduction factors, pSmad-1,5,8/9, were also reduced (Fig. 5 K, L). Thus γ1 CaMKII directs and is required for active Bmp signaling within the ventral somite region that forms sclerotome.

**Fig. 5.**
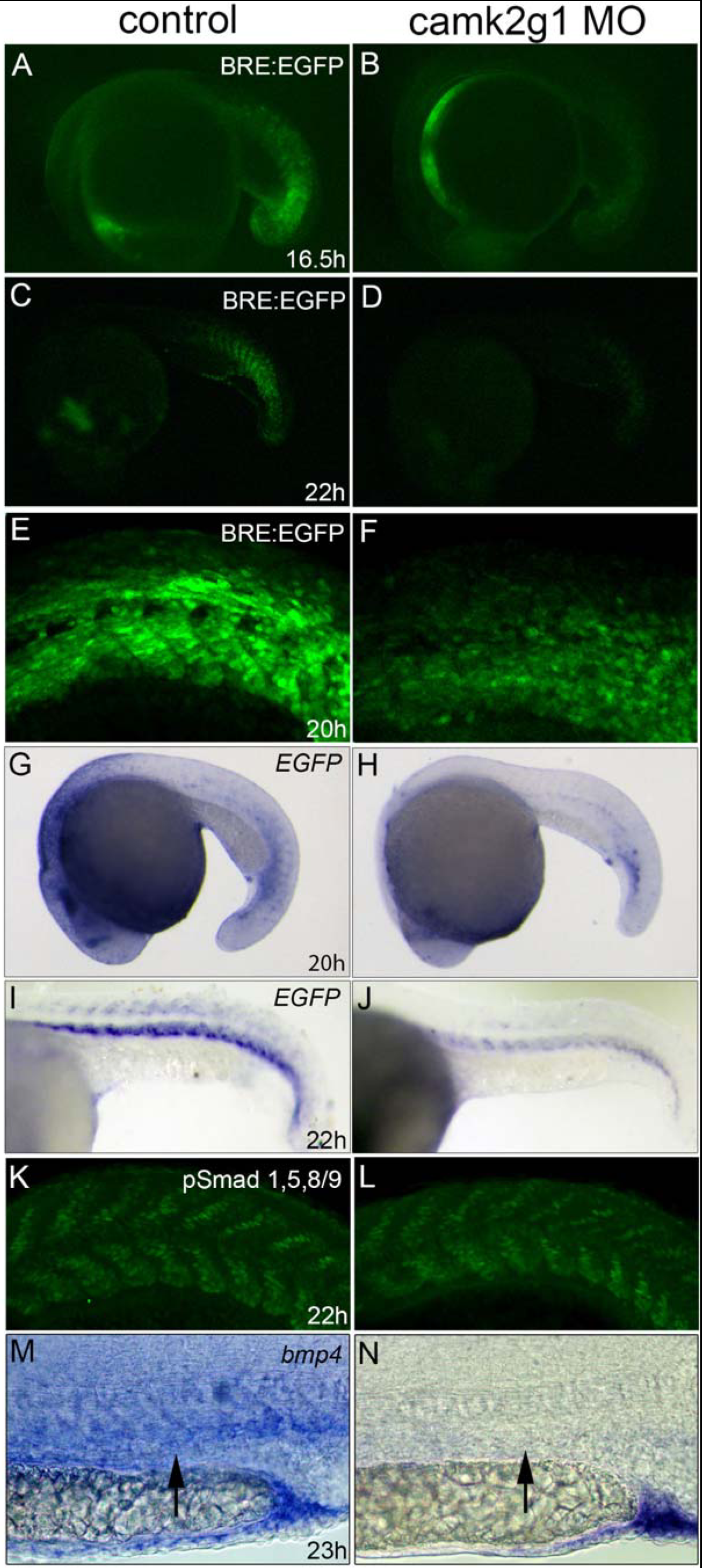
γ1 CaMKII is required for BMP signaling and expression. Uninjected *Tg(BRE:EGFP)* embryos have higher GFP fluorescence (A, C) compared to *camk2g1* morphants (B, D) at 16.5 and 22 hpf. Higher magnification views show increased fluorescence in the somites of controls (E) compared to *camk2g1* morphants (F) at 20 hpf. Consistent with reduced GFP fluorescence, *EGFP* transcript is diminished in *camk2g1* morphants compared to controls at 20 and 22 hpf (G-J). pSmad-1,5,8/9 levels are reduced in *camk2g1* morphants (L) compared to control embryos (K) at 22 hpf. *Bmp4* expression in trunk mesoderm (arrows) is severely reduced in *camk2g1* morphants (N) compared to controls (M), but displays normal expression in the medial fin and cloaca at 23 hpf. Lateral views, anterior left, dorsal up.

To better understand the basis for γ1 CaMKII regulation of Bmp signaling, we looked for changes in expression of Bmp antagonists, since their loss has been previously shown to affect sclerotome development (McMahon et al., 1998; Stafford et al., 2011; Stafford et al., 2014). *Noggin1* was expressed normally in the ventral mesoderm and *noggin2* was expressed in the notochord in *camk2g1* morphants (Fig. S4 A-D), making it unlikely that altered expression of Bmp antagonists is the explanation for altered sclerotome patterning.

During sclerotome specification, *bmp4* is expressed in ventral somite tissue and the cloaca of control embryos (Fig. 5 M). *Bmp4* expression was absent or reduced in specific regions of *camk2g1* morphants (Fig. 5 M, N) and *camk2g1* CRISPR embryos (Fig. S5 A, B), including ventral somite tissue, although it remained strongly expressed in the cloaca and medial fin at 23 hpf. Our results define γ1 CaMKII as a critical regulator of *bmp4* expression — and thus Bmp signaling — in sclerotome-fated tissues during the specification time window.

### Activated Bmp signaling alters sclerotome specification

In mouse, in the absence of secreted Bmp inhibitors (and consequently elevated BMP signaling) a select set of Shh-dependent sclerotome genes, including *Pax1* and *Pax9*, is repressed (McMahon et al., 1998; Stafford et al., 2011). To determine if ectopic Bmp signaling alters sclerotome gene expression in zebrafish, we employed a transgenic line carrying inducible expression of a constitutively active (Q207D) form of the human ALK6 (BMPR1B) BMP receptor fused to a self-cleaving photo-switchable (green-to-red) Kikume fluorophore under the control of a heat shock promoter (*hs:caALK6-KikGR*; (Row et al., 2018). Embryos were heat shocked at 40°C for 30 minutes (mins) to induce the transgene and CA-ALK6-positive embryos were identified by fluorescence after 3-4 hours. CA-ALK6-positive embryos displayed expected ectopic active Bmp signal transducers pSmad-1,5,8/9 throughout the trunk (Fig. 6 A, B), quantifiable as approximately five times more pSmad-1,5,8/9 compared with CA-ALK6 negative embryos (Fig. 6 C), indicating high activation of Bmp signaling. We verified phenotypic indicators of elevated Bmp signaling previously observed, including loss of muscle pioneer markers (Dolez et al., 2011): diminished *eng1a* expression and reduced Eng1a/Prox1 double positive muscle pioneer cells (Fig. S6 A-D). Expression of MF20, which recognizes the myosin II heavy chain, was not significantly altered, however F59 labeled slow-muscle sarcomeres appeared less organized in CA-ALK6 embryos (Fig. S6 E-H). Our results confirm this line can be used to analyze the effects of activated Bmp signaling during development.

**Fig. 6.**
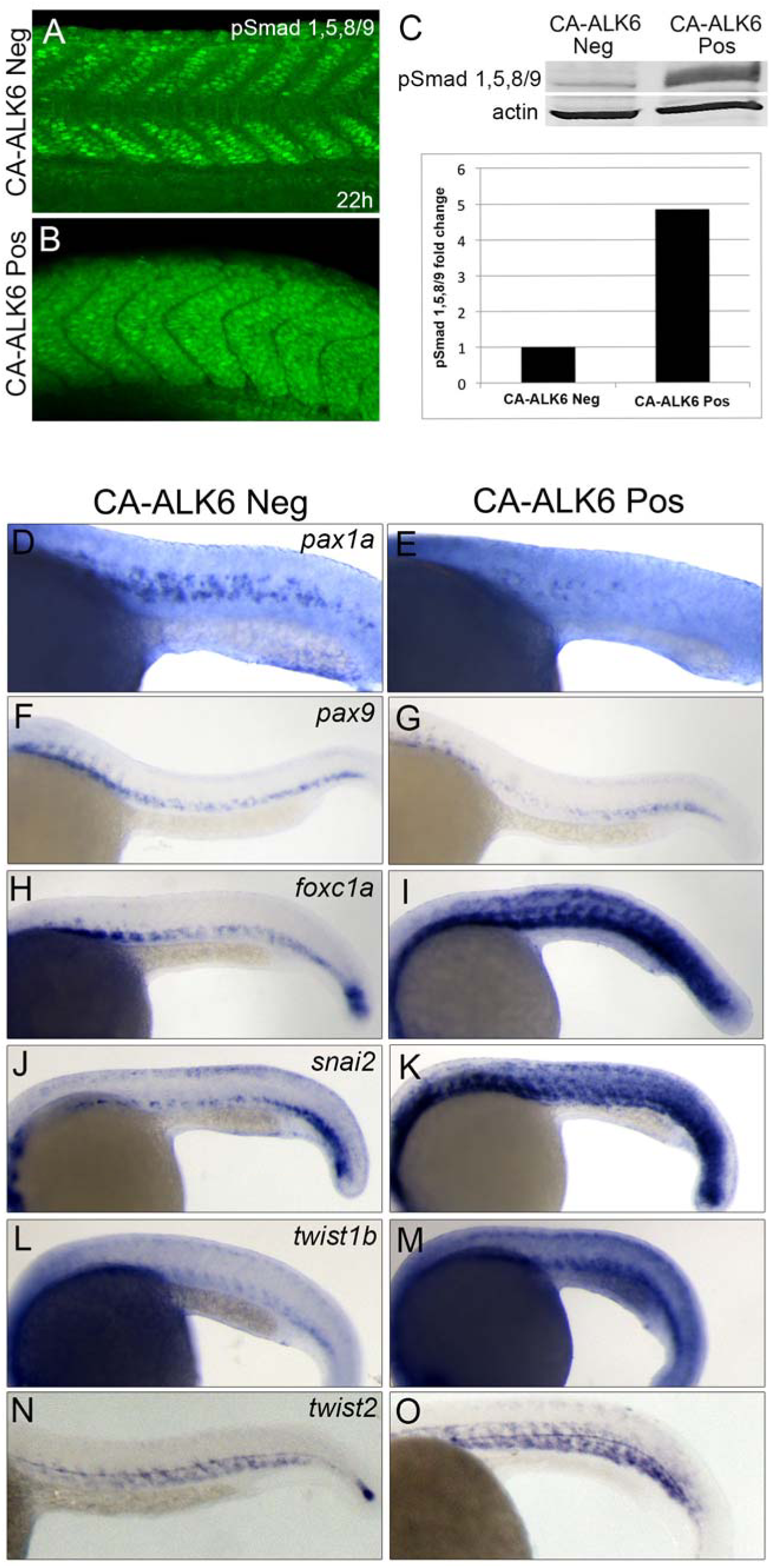
Expression of a CA-ALK6 in zebrafish embryos activates the Bmp signaling pathway and alters sclerotome gene expression. pSmad-1,5,8/9 expression is ubiquitous in CA-ALK6 positive embryos (B) compared to CA-ALK6 negative embryos at 22 hpf (A). CA-ALK6 expressing embryos have five times the expression of pSmad-1,5,8/9 compared to CA-ALK6 negative embryos at 24 hpf (C). Embryos positive for CA-ALK6 exhibit decreased expression of *pax1a* (E) and *pax9* (G) compared to CA-ALK6 negative controls (D, F). Expression of *foxc1a* (I), *snai2* (K), *twist1b* (M) and *twist2* (O) are upregulated in CA-ALK6 positive compared to negative embryos (H, J, L, N). A, B, D-O, lateral views, anterior left, dorsal up.

Elevated BMP signaling in mouse mutants carrying null alleles of BMP antagonists leads to a loss of *Pax1* and *Pax9* expression in the sclerotome (McMahon et al., 1998; Stafford et al., 2011). Zebrafish embryos expressing CA-ALK6 likewise had a reduction in *pax1a,* and *pax9,* expression (Fig. 6 D-G). However, in accord with our observation that some sclerotome genes require Bmp signaling, expression of CA-ALK6 significantly increased and expanded a distinct set of markers, including *foxc1a, snai2*, and *twist1b* throughout the somite (Fig. 6 H-M). Although *twist2* transcript abundance was elevated within the sclerotome, it was not ectopically expressed in surrounding tissues (Fig. 6 N, O). Our results demonstrate that increased Bmp signaling inhibits one set of Shh-dependent sclerotome genes, while enhancing the expression of a distinct set of Bmp-dependent markers.

### Bmp acts downstream of γ1 CaMKII

Diminished γ1 CaMKII activity led to loss of *bmp4* in the ventral somite (Fig. 5 M, N) and decreased expression of Bmp-dependent sclerotome markers (Figs. 1, 3, 6) implying that Bmp acts downstream of γ1 CaMKII to pattern sclerotome. To explicitly examine whether γ1 CaMKII acts up or downstream of Bmp signaling, we performed a rescue experiment. Wild type or *hs:caALK6-KikGR* embryos were injected with the *camk2g1* MO, heat shocked, and analyzed for expression of *foxc1a, twist1b*, and *snai2*. As observed previously, expression of CA-ALK6 expanded expression of these sclerotome genes throughout the trunk compared to control embryos (Figure 7 A-F). In addition, ectopic expression of CA-ALK6 was capable of rescuing and increasing Bmp-dependent marker expression in *camk2g1* morphants (Figure 7 G-L), indicating that Bmp signaling acts downstream of γ1 CaMKII to promote a specific subset of Bmp-dependent sclerotome genes.

**Fig. 7.**
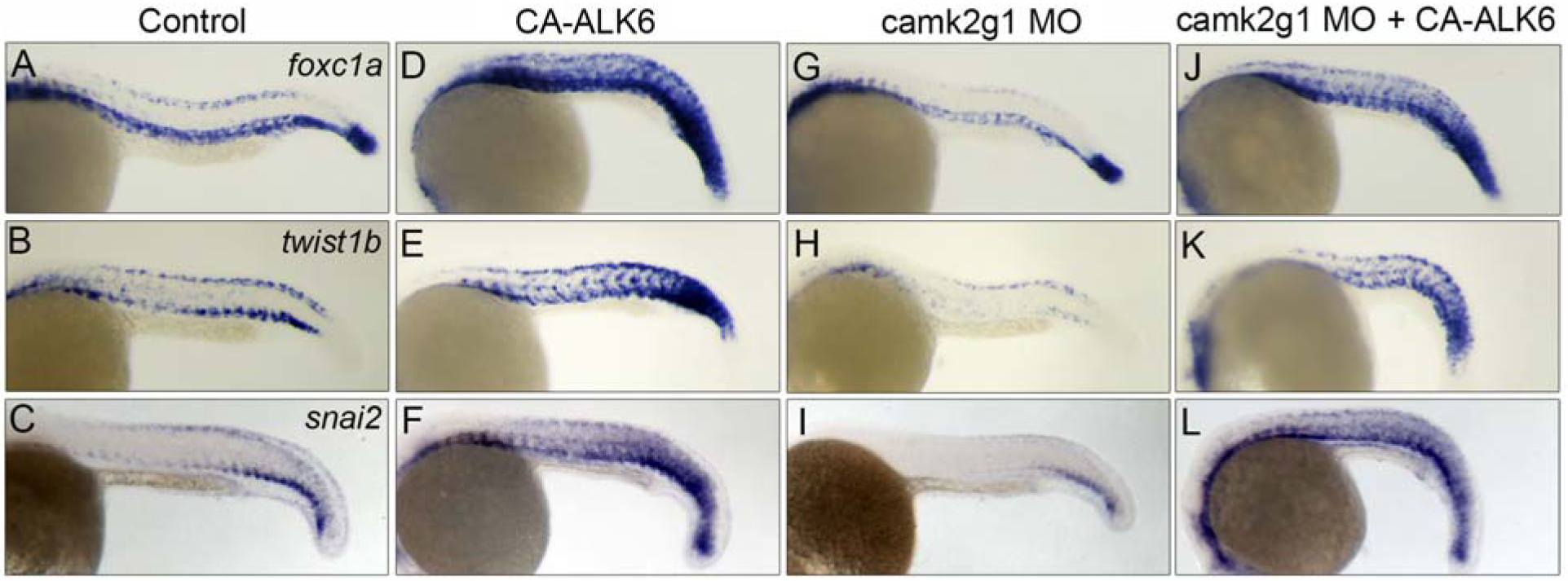
Expression of CA-ALK6 in *camk2g1* morphants rescues Bmp-dependent sclerotome gene expression. Expression of *foxc1a* (A, D, G, J), *twist1b* (B, E, H, K) and *snai2* (C, F, I, L) was analyzed in control (A-C), CA-ALK6 positive (D-F), *camk2g1* morphants (G-I), and *camk2g1* morphants expressing CA-ALK6 (J-L) at 23 hpf. Expansion of *foxc1a*, *twist1b*, and *snai2* in CA-ALK6 positive embryos (D-F), compared to controls (A-C) can overcome the reduction of these markers in *camk2g1* morphants (G-I vs. J-L). Lateral views, anterior left, dorsal up.

## Discussion

Our data reveal γ1 CaMKII as a pivotal regulator of sclerotome specification and compartmentalization, upstream of Shh and Bmp signaling. Sclerotome comprises a migratory population of cells with highly diverse adult tissue fates including vertebrae, vertebral discs, ribs, sternum, tendons, myotendinous junctions, vascular smooth muscle, and other tissues (Christ et al., 2004; Monsoro-Burq et al., 1996; Stickney et al., 2000). How early do sclerotome cells achieve distinct gene signatures? How and when are the specific genetic programs differentially engaged to drive discrete fate potentials? Are cells with certain fate potentials predisposed to migrate to unique sites of adult differentiation?

A careful analysis of sclerotome markers over time reveals that there is dynamic and spatially non-uniform gene expression, suggesting the presence of early subcompartments (Christ et al., 2004; Hughes et al., 2009; Monsoro-Burq, 2005), and indicates that there must be regulatory control of differential gene expression within the sclerotome. Specific combinatorial gene signatures might reflect discrete migratory capabilities and fate potentials. A number of potential compartments have been previously proposed, correlating to adult tissue contributions, including arthrotome, syndetome, meningotome, and neurotome (Christ et al., 2004), but little is known about gene signatures that might portend these fates, or how and when they are instructed.

From work in mouse and chick, SHH signaling is a well characterized essential signal directing both formation of sclerotome (Chiang et al., 1996; Fan and Tessier-Lavigne, 1994; Johnson et al., 1994; Teillet et al., 1998) and differential fates within the overall somite by acting as a morphogen (Cairns et al., 2008). One level of control over sclerotome compartmentalization could come from differential distribution of SHH and distinct genetic threshold responses. Our data define a number of clear sclerotomal targets of zebrafish Shh signaling exhibiting diminished or lost expression in cyclopamine-treated embryos that are consistent with prior studies, including *pax1a*, *pax9*, and *twist2* (Arnold et al., 2015; Ma et al., 2018). Our data reveal that γ1 CaMKII regulates shaping of this morphogen gradient, by promoting secretion and/or diffusion of Shh. Consequently, in *camk2g1* morphants, we see loss of this set of Shh-dependent sclerotome target genes. Interestingly, a second set of sclerotome genes is independent of Shh signaling, including *foxc1a*, *twist1b*, and *snai2*, which despite being regulated by γ1 CaMKII, show little or no diminution when Shh is inhibited. These results imply the existence of a second parallel pathway downstream of γ1 CaMKII that instructs sclerotome specification.

Bmp is that secondary signaling pathway responsible for activating expression of a distinct set of sclerotome genes. Previous work in mouse had shown that limiting BMP signaling through expression of secreted BMP inhibitory factors, including noggin and follistatin, is essential for permitting expression of sclerotome markers including *Pax1* and *Pax9* (McMahon et al., 1998; Stafford et al., 2011; Stafford et al., 2014). Our data confirm that Bmp signaling in zebrafish does indeed limit expression of *pax1a* and *pax9*, and that too much Bmp signaling leads to loss of these markers. However, we observe that a distinct set of sclerotome markers, including *foxc1a*, *twist1b*, and *snai2* are activated by Bmp signaling and rely on it for their expression.

Interestingly, our data show that Bmp-dependent Smads are most active in the anterior half of somites (Fig. 5K), in agreement with previous studies (Maurya et al., 2011; Wilkinson et al., 2009). The putative Bmp targets, *foxc1a*, *twist1b*, and *snai2*, are preferentially expressed in the anterior half of the sclerotomes (Fig. 2 G, I, K), whereas genes inhibited by Bmp signaling, *pax1a*, *pax9*, and *twist2*, are expressed in the posterior part of the sclerotome (Fig. 1, I, K; Fig. 2 E). Together, our data support a model where Shh drives posterior and Bmp drives anterior sclerotomal gene expression, as parallel signaling arms downstream of γ1 CaMKII. It is also likely that signaling is limited and gene expression is further modified by secreted Bmp inhibitors (McMahon et al., 1998; Stafford et al., 2011; Stafford et al., 2014), as well as overlayed signaling pathways such as fibroblast growth factor and Notch (Koudijs et al., 2008), and feedback inhibition (this study; (Koudijs et al., 2008), leading to dynamic gene expression over time and stabilization of compartmentalized gene signatures. Our model provides a conceptual basis for the formation of distinct sclerotome subdomains exhibiting specific combinations of genes based on inductive, permissive, and inhibitory signaling. Future research will assess the specific fate potentials driven by particular combinations of genes, but presumably they control of specification, lineage restriction, and migration to the site of adult residency.

## Materials and Methods

### Zebrafish strains and care

Wild type (AB), *Tg(hsp70l:dnXla.Bmpr1a-GFP)^w30^* (referred to as *hs:dnBmpr-GFP* in the text; Pyati et al., 2005), *Tg(BMPRE-AAV.Mlp:EGFP)^mw29^* (referred to as *BRE:EGFP* in the text; Collery and Link, 2011), and *Tg(hsp70l:caHsa.ALK6-2A-NLS-KikGR)^sbu106^*(referred to as *hs:caALK6-KikGR* in the text; Row et al., 2018) embryos were obtained through natural spawnings and raised at 28.5°C as previously described (Kimmel et al., 1995; Laux et al., 2011; Pyati et al., 2005) under an approved IACUC protocol, and in compliance with all IACUC and AVMA guidelines. Embryos were not preferentially selected by sex.

### Heat Shock Lines

*Hs:caALK6-KikGR* embryos were heat shocked at 40°C for 30 minutes (mins) and allowed to develop for 3-4 hours (hrs). Kikume positive and negative embryos were identified by fluorescence and processed for analysis. *Hs:dnBmpr-GFP* embryos were heat shocked at 37°C for 20 mins and allowed to develop for 3 hrs. GFP positive and negative embryos were identified and processed for analysis.

### Fluorescence Localization

Embryos were fixed in 4% formaldehyde and incubated with rabbit anti-phospho-Smad-1,5,8/9 (Cell Signaling, 13820, 1:20), rabbit anti-γ1/δ2 CaMKII (GeneTex, GTX128842, 1:20), rabbit anti-Shha (AnaSpec,, AS-55574, 1:20), rabbit anti-human PROX1 (ReliaTech, 1:20), mouse anti-engrailed (Developmental Studies Hybridoma Bank, University of Iowa (4D9) 1:100), mouse anti-myosin, sarcomere (Developmental Studies Hybridoma Bank, University of Iowa (MF20) 1:100), or mouse anti-myosin heavy chain, slow muscle (Developmental Studies Hybridoma Bank, University of Iowa (F59) 1:100). Primary antibody incubations were overnight and were visualized with secondary of either goat anti-mouse Alexa 488, goat anti-rabbit Alexa 488, goat anti-mouse Alexa 568, or goat anti-rabbit Alexa 568 (Invitrogen at 2.5 mg/ml) using confocal microscopy (Nikon C1 Plus two-laser, Nikon C2 Plus four-laser) on a Nikon E-600 compound microscope using a 20X dry or 40x dry objective, or on Nikon AZ-100 macro zoom fluorescent stereo microscope.

### Morpholino and mRNA injections

A previously described translation-blocking antisense morpholino oligonucleotide (MO) targeted against zebrafish *camk2g1* (1 ng) was injected at the 1-4 cell stage (Francescatto et al., 2010; Rothschild et al., 2011; Rothschild et al., 2013).

### Camk2g1 CRISPR/Cas9 G0 embryos

Four guide RNAs (gRNAs) against *camk2g1* were designed and generated as previously described (Rothschild, et. al., 2020). The four gRNAs were co-injected at 200ng/μl each with Cas9 protein (1ng; New England Biosciences) into one-cell stage zebrafish embryos.

### Whole mount *in situ* hybridization

Digoxigenin-labeled anti-sense riboprobes (0.5–1.5 kb) were synthesized using T3, T7, or SP6 RNA polymerase from cloned cDNAs and then hybridized with fixed embryos as previously described (Clements et al., 2011; Rothschild et al., 2007). Synthesis of *pax1a*, *twist2*, *foxc1a*, *twist1b*, *snai2*, *shha*, *bmp4*, and *EGFP* probes and parent constructs were described previously (Clements et al., 2011). Probes *nog1* (S 5’-AGGATGGATTTCCCGCGGTGTTTG-3’, AS 5’-GGTCAGTTCGCGCATGAGCATTTG-3’), *nog2* (S 5’-CATGGGCAGCATCACCCGCGCGCT-3’, AS 5’-CTCAGCACGAGCACTTGCACTGGG-3’), and *pax9* (F 5’-tctagaATGGAGCCAGCCTTT-3’, R 5’-atggatccTCATAGAGCTGAAGCCACCAG-3’) were generated by PCR amplification from 24 hpf whole embryo cDNA and cloned to pCRII-TOPO (Invitrogen) according to the manufacturer’s instructions. For whole mount *in situ* hybridization, embryos were developed using alkaline phosphatase-conjugated anti-digoxigenin.

### Drug Treatment

Zebrafish embryos were treated with 10 μM DMH1 (Sigma Aldrich) or 100 μM cyclopamine (Cayman Chemical) for 2-4 hrs in 1 ml E3 embryo media. Embryos were dechorionated prior to drug treatment.

### Western blots

Embryos were manually dechorionated, transferred to cold Ringer’s solution with 1 mM EDTA and 0.3 mM phenylmethylsulfonylfluoride (PMSF), and triturated to remove yolk (Westerfield, 1993). Embryos were washed three times in cold Ringer’s solution and immediately placed in Laemmli Buffer with DTT. Equivalent amounts of soluble protein were immunoblotted as described (Lantsman and Tombes, 2005) using a rabbit anti-phospho-Smad-1,5,8/9 antibody (Cell Signaling) or a mouse anti-actin (Sigma) antibody (Rothschild et al., 2009).

## Acknowledgments

The authors wish to acknowledge Clair M. Kelley for the *pax9* probe. This work was supported by ALSAC funding, NHLBI R00HL097150, NIDDK R01DK113973, and March of Dimes #5-FY14-42 awards to W.K.C.; a Massey Cancer Center Pilot Project Grant to S.C.R.; and NSF IOS 1452928, American Heart Association 13SDG14360032, and NIGMS R01GM124282 awards to B.L.M.

## Author Contributions

S.C.R. and W.K.C. conceived the experiments, wrote the manuscript, analyzed results, and secured funding; S.C.R. performed the experiments; R.H.R. and B.L.M. generated and shared the *Tg(hsp70l:caHsa.ALK6-2A-NLS-KikGR)^sbu106^* line prior to publication; B.L.M. and R.H.R. provided critical comments on the manuscript.

## Declaration of Interests

The authors declare no competing interests.

---

**Fig. S1.**
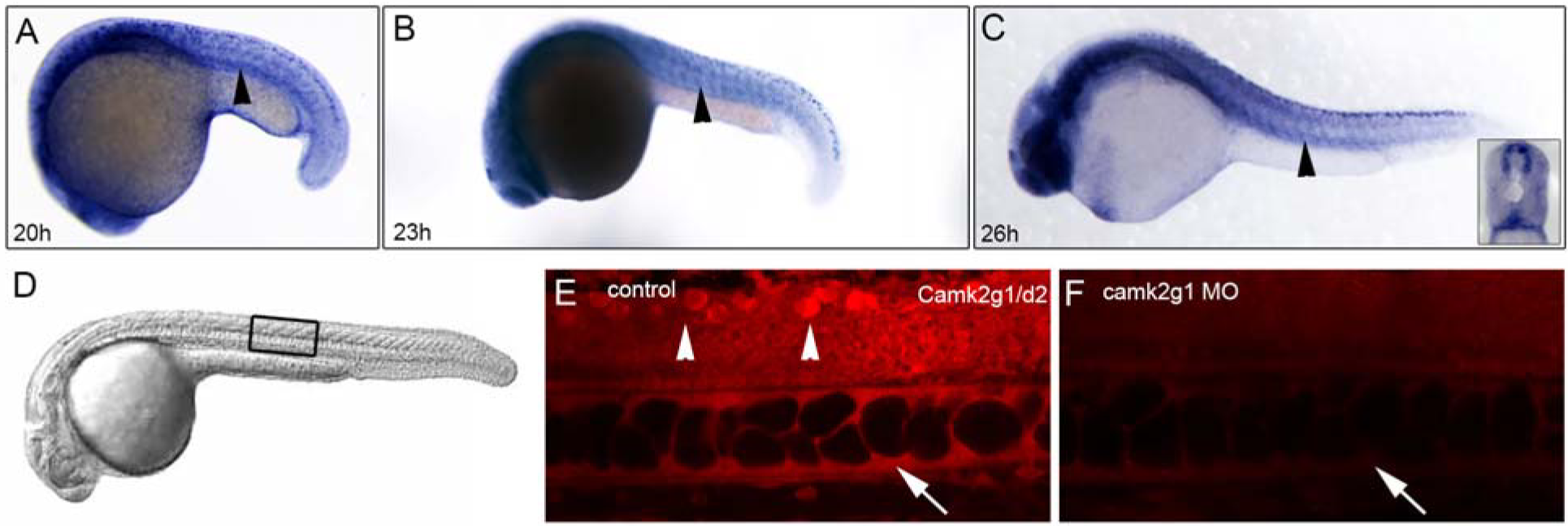
*Camk2g1* is expressed in the sclerotome during zebrafish development. *Camk2g1* mRNA is expressed in ventromedial somite tissue where sclerotome is located at 20, 23, and 26 hpf (A-C, arrowheads) as well as in neural tube, spinal cord cell bodies, brain, and notochord (A-C). An antibody recognizing both δ2 CaMKII and γ1 CaMKII confirms expression in the notochord (E, arrow), neural tube and spinal cord cell bodies (E, arrowheads) consistent with the location of *camk2g1*, but not *camk2d2* mRNA expression. Morpholino targeting of *camk2g1* leads to loss of all CaMKII signal (E vs. F) in the notochord (arrows), spinal cord, and spinal cord cell bodies (arrowheads). Lateral views, anterior left, dorsal up.

**Fig. S2.**
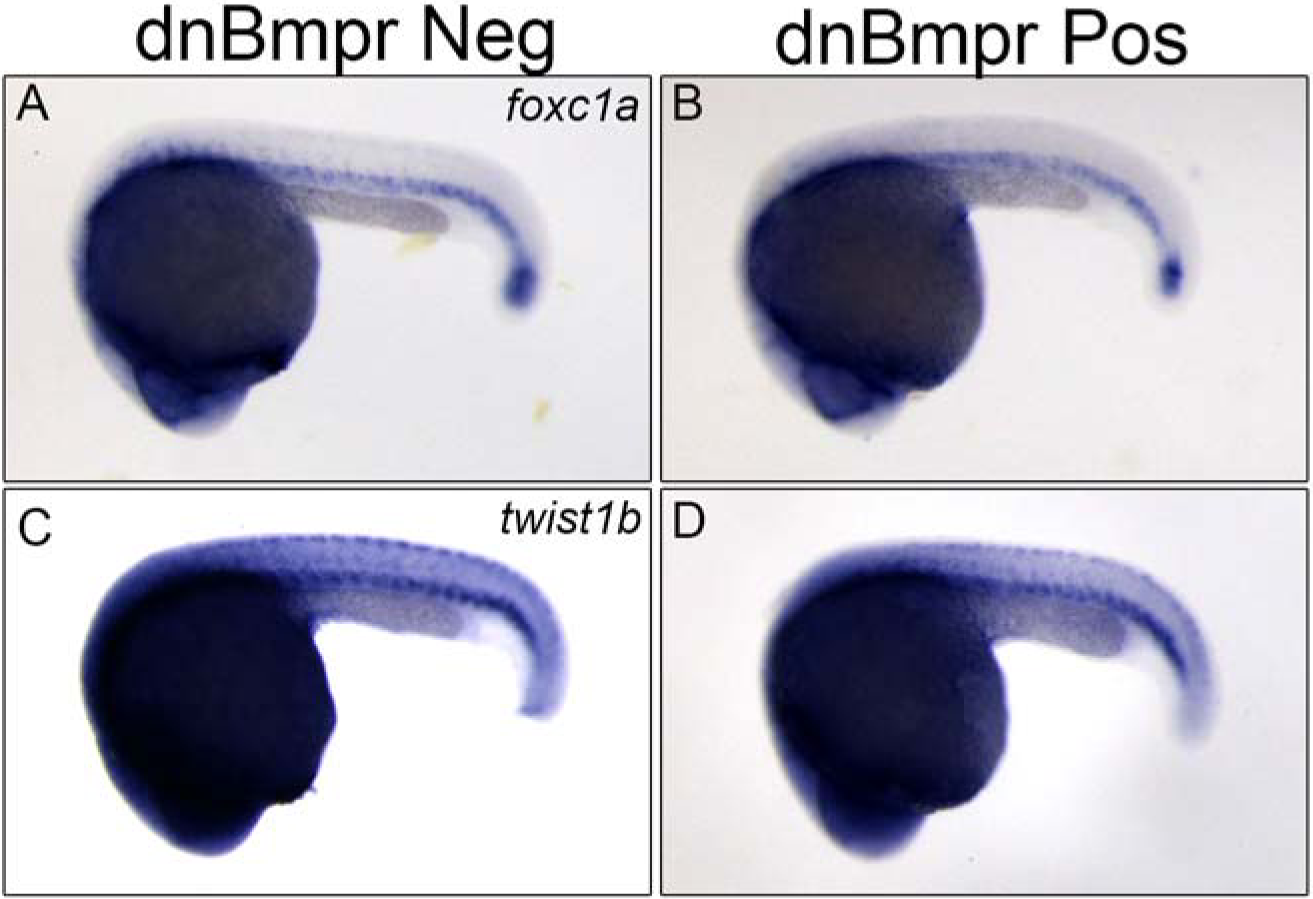
Inhibition of Bmp signaling reduces expression of Bmp-dependent sclerotome genes. Expression of *foxc1a* (B) and *twist1b* (D) is reduced in dnBmpr positive embryos compared to embryos negative for dnBmpr (A, C). Lateral views, anterior left, dorsal up.

**Fig. S3.**
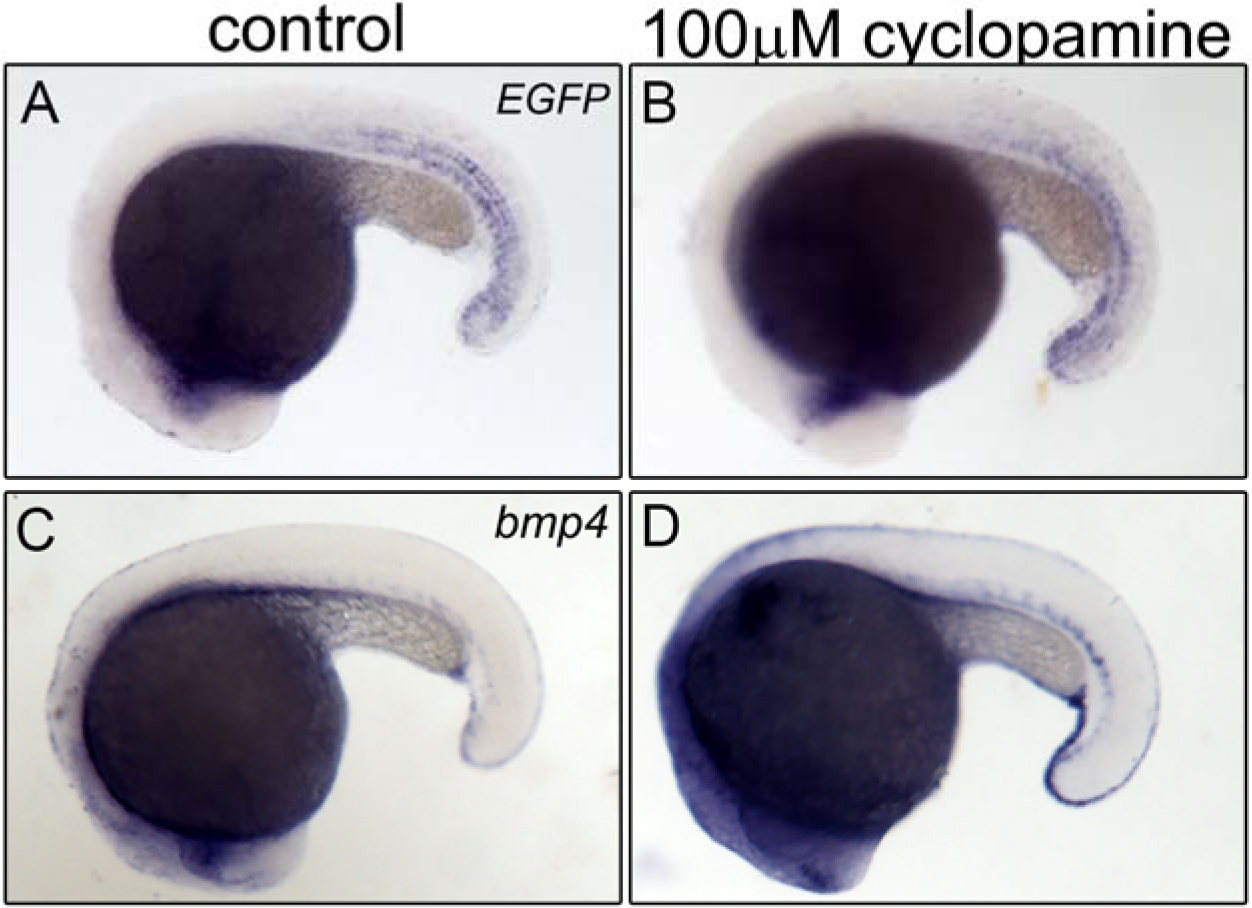
Effect of Shh inhibition on Bmp signaling. Treatment of embryos with 100 μM cyclopamine does not alter *EGFP* message in *BRE:EGFP* fish (B) compared to control embryos (A). Expression of *bmp4* in the ventral mesoderm slightly increases in cyclopamine treated embryos (D) compared to control embryos (C). 18 hpf, lateral views, anterior left, dorsal up.

**Fig. S4.**
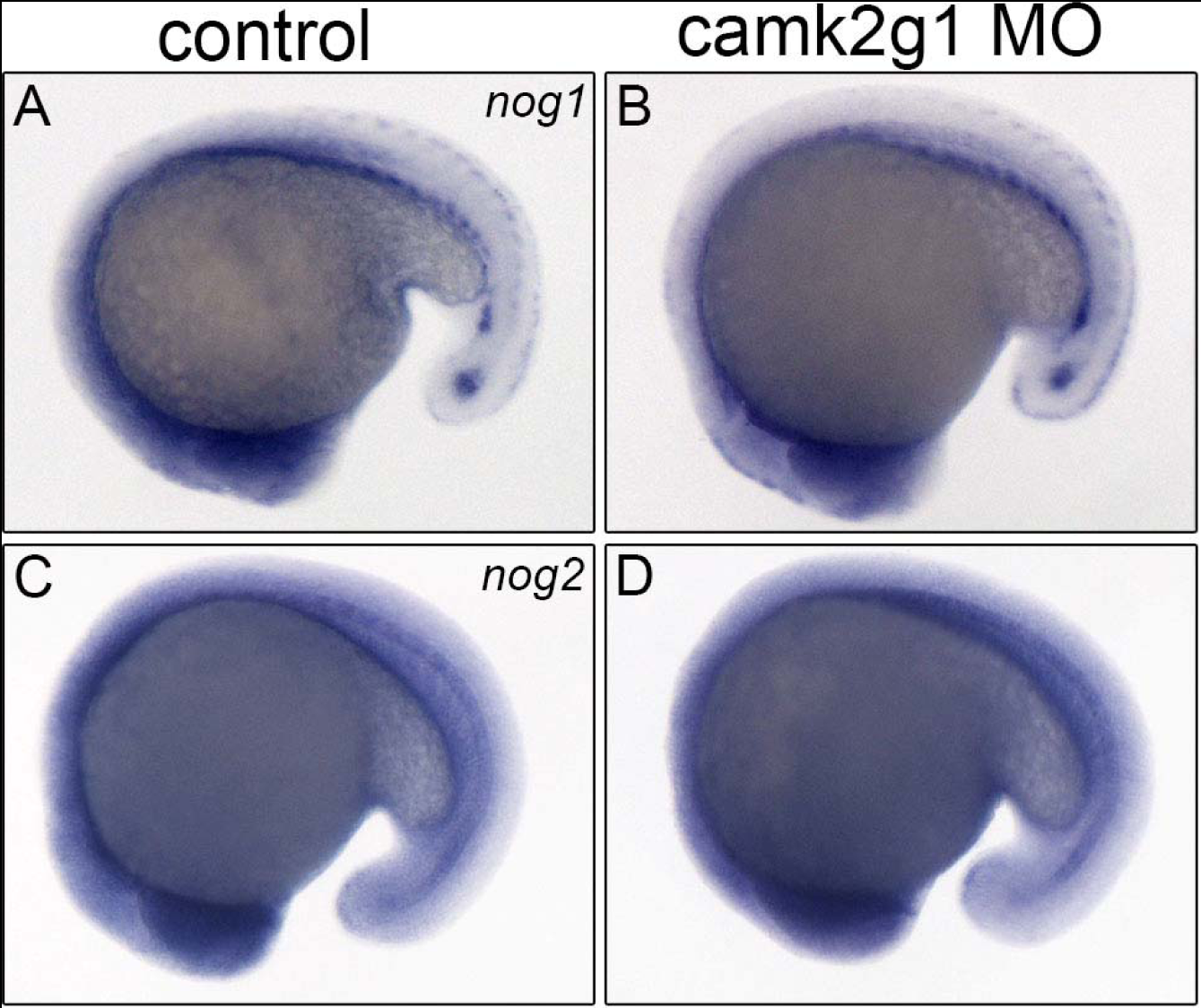
Normal expression of Bmp antagonists in *camk2g1* morphants. Expression of *nog1* in the ventral mesoderm(A, B) and *nog2* in the notochord (C, D) are equivalent to control embryos at 18 hpf. Lateral views, anterior left, dorsal up.

**Fig. S5.**
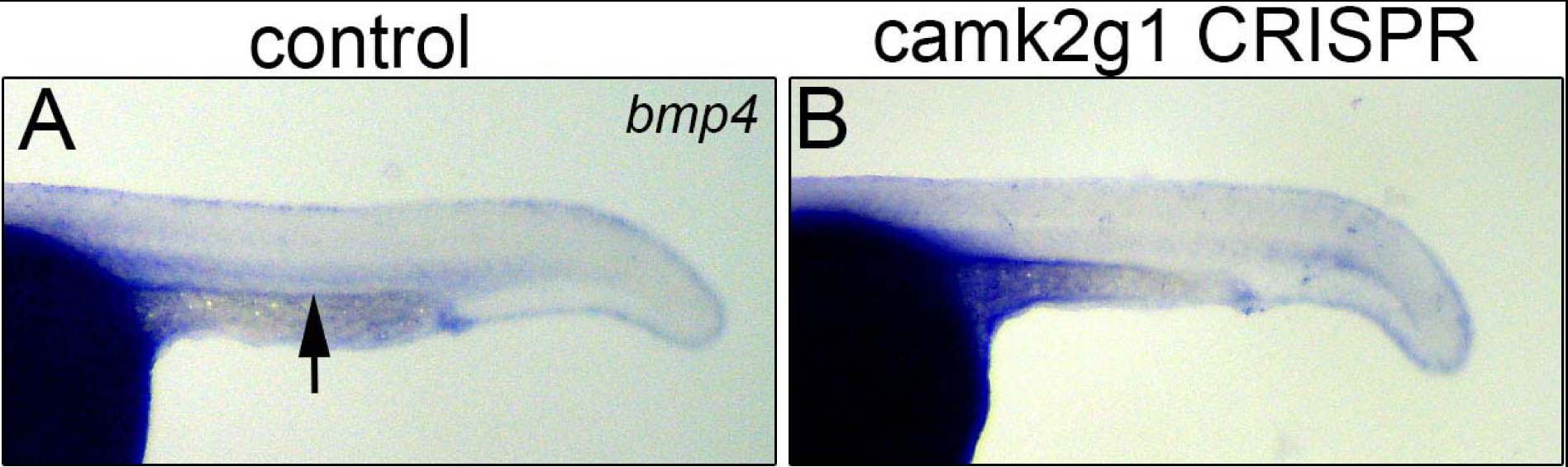
Reduced *bmp4* expression in *camk2g1* CRISPR embryos. Expression of *bmp4* in the ventral mesoderm is reduced in *camk2g1* CRISPR embryos (B) compared to control embryos at 24 hpf (A). Lateral views, anterior left, dorsal up.

**Fig. S6.**
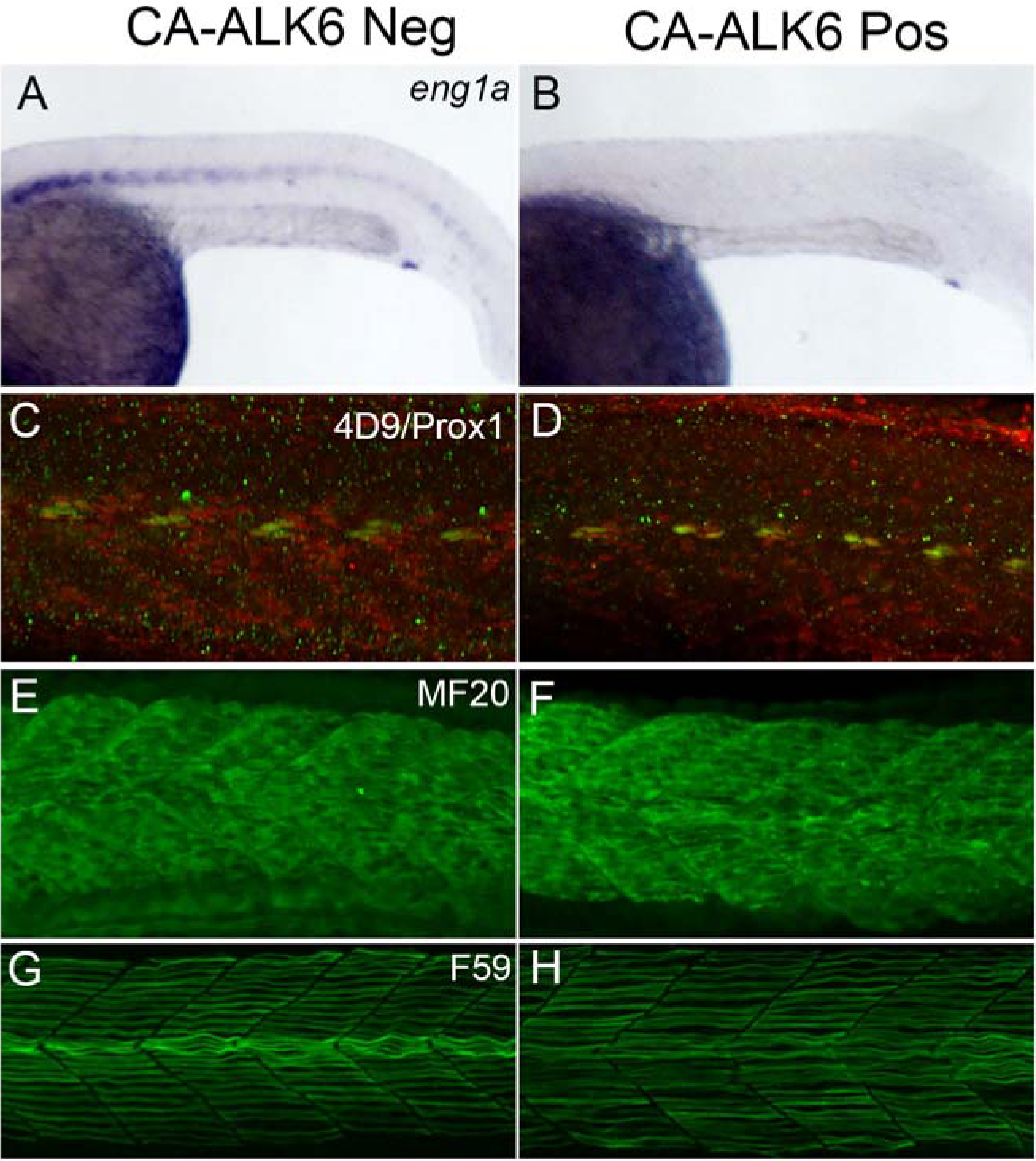
*Eng1a* expression is reduced in CA-ALK6 embryos. *Eng1a* is expressed in muscle pioneer cells in CA-ALK6 negative embryos (A) and absent in CA-ALK6 positive embryos (B). 4D9 Prox1 double positive muscle pioneer cells are reduced in CA-ALK6 positive embryos (D) compared to CA-ALK6 negative embryos (C). Fast muscle (MF20) is unaffected in CA-ALK6 negative and positive embryos (E, F). Slow muscle (F59) appears disorganized in CA-ALK6 positive embryos (H) compared to CA-ALK6 negative embryos (G). Lateral views, anterior left, dorsal up.

**Table S1:**

| Type of embryo | Gene Tested | Stage | Down | No Change | Up | Figure | Description |
| --- | --- | --- | --- | --- | --- | --- | --- |
| camk2g1 MO | foxc1a | 24hpf | 43 | 5 | 0 | Figure 1B |  |
| camk2g1 CRISPR | foxc1a | 24hpf | 25 | 4 | 0 | Figure 1C |  |
| camk2g1 MO | twist1b | 24hpf | 22 | 5 | 0 | Figure 1E |  |
| camk2g1 CRISPR | twist1b | 24hpf | 18 | 8 | 0 | Figure 1F |  |
| camk2g1 MO | twist2 | 24hpf | 24 | 3 | 0 | Figure 1H |  |
| camk2g1 CRISPR | twist2 | 24hpf | 23 | 3 | 0 | Figure 1I |  |
| camk2g1 MO | snai2 | 24hpf | 18 | 10 | 0 | Figure 1K |  |
| camk2g1 CRISPR | snai2 | 24hpf | 24 | 4 | 0 | Figure 1L |  |
| camk2g1 MO | pax1a | 24hpf | 16 | 4 | 0 | Figure 1N |  |
| camk2g1 CRISPR | pax1a | 24hpf | 22 | 6 | 0 | Figure 1O |  |
| camk2g1 MO | pax9 | 24hpf | 18 | 2 | 0 | Figure 1Q |  |
| camk2g1 CRISPR | pax9 | 24hpf | 14 | 6 | 0 | Figure 1R |  |
| 100µM cyclopamine | pax1a | 20hpf | 21 | 4 | 0 | Figure 2B |  |
| 100µM cyclopamine | pax9 | 20hpf | 26 | 3 | 0 | Figure 2D |  |
| 100µM cyclopamine | twist2 | 20hpf | 21 | 12 | 0 | Figure 2E |  |
| 100µM cyclopamine | foxc1a | 22hpf | 0 | 26 | 0 | Figure 2H |  |
| 100µM cyclopamine | twist1b | 22hpf | 0 | 25 | 0 | Figure 2J |  |
| 100µM cyclopamine | snai2 | 20hpf | 0 | 21 | 0 | Figure 2L |  |
| 10µM DMHI | foxc1a | 20hpf | 32 | 10 | 0 | Figure 3B |  |
| 10µM DMHI | twist1b | 20hpf | 18 | 8 | 0 | Figure 3D |  |
| 10µM DMHI | snai2 | 20hpf | 22 | 6 | 0 | Figure 3F |  |
| 10µM DMHI | pax1a | 20hpf | 0 | 24 | 0 | Figure 3H |  |
| 10µM DMHI | pax9 | 18hpf | 0 | 31 | 0 | Figure 3J |  |
| 10µM DMHI | twist2 | 18hpf | 2 | 22 | 0 | Figure 3L |  |
| camk2g1 MO | shha | 14hpf | 0 | 18 | 0 | Figure 4F |  |
| camk2g1 MO | shha | 18hpf | 0 | 10 | 31 | Figure 4H |  |
| camk2g1 MO | shha | 24hpf | 0 | 12 | 23 | Figure 4J |  |
| camk2g1 MO | BRE:EGFP | 15ss | 67 | 6 | 0 | Figure 5B |  |
| camk2g1 MO | BRE:EGFP | 22hpf | 56 | 12 | 0 | Figure 5D |  |
| camk2g1 MO | BRE:EGFP | 20hpf | 12 | 4 | 0 | Figure 5F |  |
| camk2g1 MO | egfp | 20hpf | 20 | 2 | 0 | Figure 5H |  |
| camk2g1 MO | egfp | 22hpf | 24 | 1 | 0 | Figure 5J |  |
| camk2g1 MO | bmp4 | 23hpf | 28 | 10 | 0 | Figure 5L |  |
| CA Alk6 positive | pax1a | 23hpf | 49 | 0 | 0 | Figure 7B |  |
| CA Alk6 positive | pax9 | 24hpf | 38 | 7 | 0 | Figure 7D |  |
| CA Alk6 positive | foxc1a | 23hpf | 0 | 0 | 35 | Figure 7F |  |
| CA Alk6 positive | snai2 | 23hpf | 0 | 0 | 20 | Figure 7H |  |
| CA Alk6 positive | twist1b | 22hpf | 0 | 0 | 32 | Figure 7J |  |
| CA Alk6 positive | twist2 | 24hpf | 0 | 3 | 28 | Figure 7L |  |
| CA Alk6 positive | foxc1a | 24hpf | 0 | 0 | 20 | Figure 8B |  |
| camk2g1 MO | foxc1a | 24hpf | 21 | 3 | 0 | Figure 8C |  |
| camk2g1 MO | foxc1a | 24hpf | 0 | 0 | 20 | Figure 8D | CA Alk6 positive |
| CA Alk6 positive | twist1b | 24hpf | 0 | 0 | 28 | Figure 8F |  |
| camk2g1 MO | twist1b | 24hpf | 18 | 3 | 0 | Figure 8G |  |
| camk2g1 MO | twist1b | 24hpf | 0 | 0 | 24 | Figure 8H | CA Alk6 positive |
| CA Alk6 positive | snai2 | 24hpf | 0 | 0 | 31 | Figure 8J |  |
| camk2g1 MO | snai2 | 24hpf | 24 | 8 | 0 | Figure 8K |  |
| camk2g1 MO | snai2 | 24hpf | 0 | 3 | 27 | Figure 8L | CA Alk6 positive |

## Notes

### Competing Interest Statement

The authors have declared no competing interest.

